# GATA3 controls mitochondrial biogenesis in primary human CD4^+^ T cells during DNA damage

**DOI:** 10.1101/727479

**Authors:** Lauren A. Callender, Johannes Schroth, Elizabeth C. Carroll, Lisa E.L. Romano, Eleanor Hendy, Audrey Kelly, Paul Lavender, Arne N. Akbar, J. Paul Chapple, Sian M. Henson

**Author notes:** Correspondence: Dr. Sian M. Henson. Both authors contributed equally (Lauren A. Callender, Johannes Schroth).

## Abstract

GATA binding protein 3 (GATA3) has traditionally been regarded as a lineage-specific transcription factor that drives the differentiation of CD4^+^ T helper (Th) 2 cells. However, increasing evidence shows that GATA3 is involved in a myriad of processes such as immune regulation, proliferation and maintenance in other T cell and non-T cell lineages. We show here a previously unknown mechanism utilized by CD4^+^ T cells to increase mitochondrial mass in response to DNA damage through the binding of GATA3, AMP-activated protein kinase (AMPK), peroxisome-proliferator-activated receptor γ co-activator-1α (PGC1α), nuclear factor erythroid 2-related factor 2 (NRF2) and superoxide dismutase 3 (SOD3) to the DNA damage repair (DDR) component ATR. These findings extend the pleotropic nature of GATA3 and highlight the potential for GATA3-targeted cell manipulation for clinical interventions.

## Main text

The function of GATA3 in the development of T cells has been well characterised. It is the only member of the GATA family expressed in the T cell lineage^1^, and during early T cell development in the thymus GATA3 is essential for CD4^+^ lineage commitment^2^. Among mature, peripheral CD4^+^ T cells, GATA3 is also required for Th2 differentiation^3,4^. More recently, GATA3 has been shown to have many functional roles beyond simply controlling CD4^+^ Th2 cell differentiation. These roles include the development of invariant NKT (iNKT) cells^5^, the maturation and homing of natural killer (NK) cells^6^ and the regulation and activation of CD8^+^ T cells^7^. Collectively these studies expand the role of GATA3 in non-T cell and CD8^+^ T cell lineages, however the role of GATA3 in non-Th2 CD4^+^ T cells is unknown.

As CD4^+^ T cells become more highly differentiated they develop a Th1-like phenotype, characterised by increased IFN-ƴ and decreased IL-4, IL-5 and IL-13 cytokine production^8,9^. Here, we examined the expression of GATA3 in primary human CD4^+^ T cell subsets defined by CD45RA and CD27 expression (Extended data figure 1a) – Naive (CD45RA^+^CD27^+^), central memory (CM; CD45RA^-^CD27^+^), effector memory (EM; CD45RA^-^CD27^-^) and effector memory that re-express CD45RA (EMRA; CD45RA^+^CD27^-^)^10,11^. CD4^+^ EMRA T cells are a highly differentiated population that exhibit many characteristics of cellular senescence such as DNA damage^12^, and have Th1-like phenotype^8,9^. Despite this, unstimulated CD4^+^ EMRA T cells expressed the highest levels of GATA3 when compared to the other three subsets by flow cytometry and qPCR (Figure 1a and extended data figure 1b). Furthermore, the expression of GATA3 remained high in the CD4^+^ EMRA subset following stimulation (Extended data figure 1c). This observation led us to hypothesize that GATA3 has an alternate function in the CD4^+^ EMRA subset. In the CD8^+^ lineage, GATA3 expression was low and remained unchanged amongst the four subsets (Extended data figure 1d).

**Figure 1:**
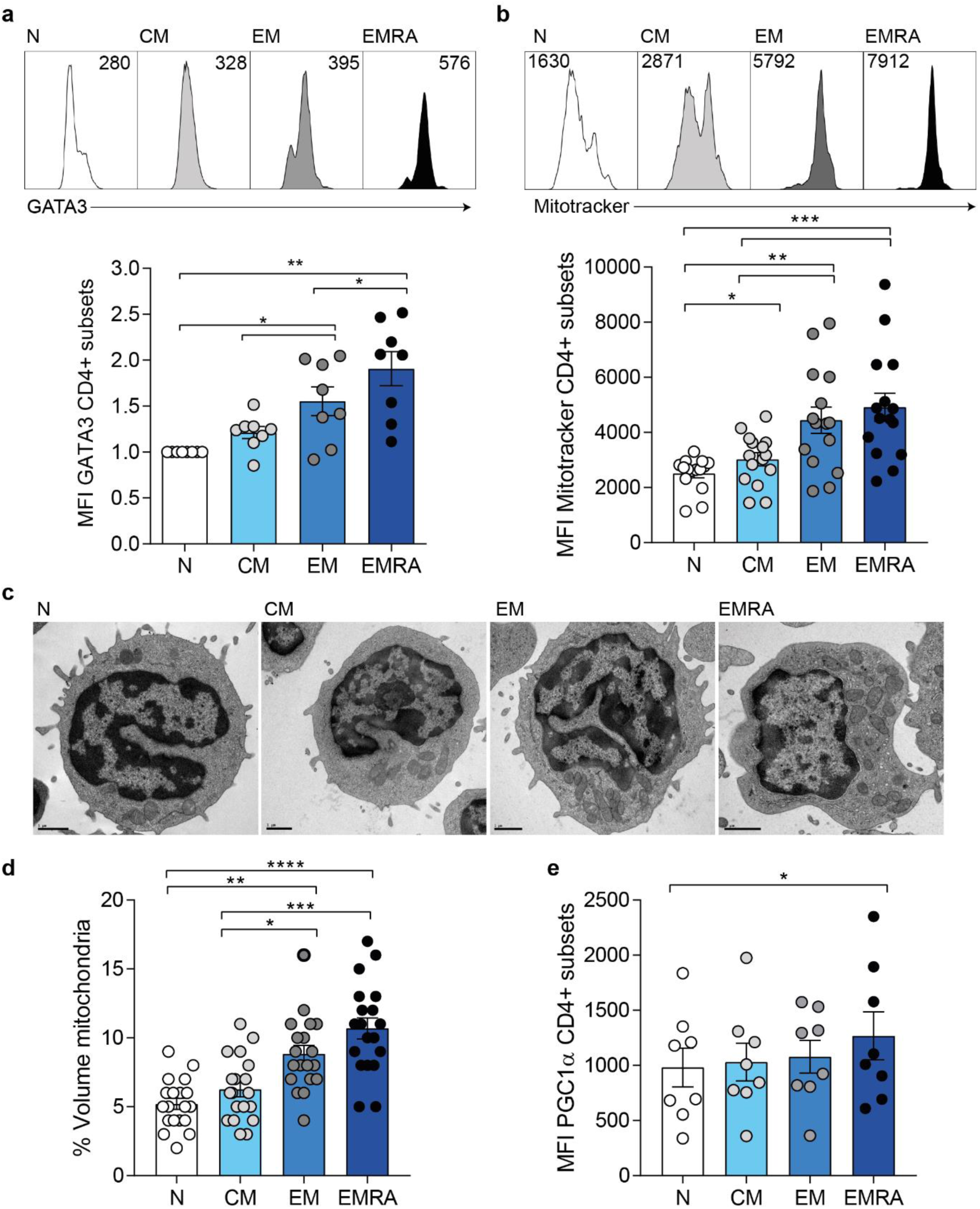
Senescent CD4^+^ EMRA T cells expressed high levels of GATA3 which correlates with increased mitochondrial mass. (a) Representative flow cytometry histograms and cumulative graph of GATA3 staining in CD27/CD45RA defined CD4^+^ T cell subsets (n=7). (b) Representative flow cytometry histograms and cumulative graph of Mitotraker Green staining in CD27/CD45RA defined CD4^+^ T cell subsets (n=14). (c) Electron microscope images of CD27/CD45RA defined CD4^+^ T cell subsets imaged directly *ex vivo*. Scale bars 1 μM. (d) Graph shows the percentage by cell volume of mitochondria in CD27/CD45RA defined CD4^+^ T cell subsets determined by a point counting grid method from 20 different electron microscope images. (e) PGC1α staining in CD45RA/CD27 defined CD4^+^ T cell subsets (n=8).

Cellular metabolism is a crucial regulator of T cell function and fate, and the physiological importance of mitochondria in regard to cell metabolism has been widely appreciated for many years. We have previously shown mitochondrial mass to be lower in CD8^+^ EMRAs compared to other memory subsets and the mitochondria present to be dysfunctional^13^. Therefore, we sought to investigate the mitochondrial properties of CD4^+^ T cells subsets. First, we assessed the total mitochondrial mass of CD4^+^ T cell subsets using the mitochondrial probe; Mitotracker Green. These data revealed that mitochondrial mass increased with differentiation in unstimulated and stimulated CD4^+^ T cells, with CD4^+^ EMRA T cells displaying the highest mitochondrial content in both cases (Figure 1b, extended data figure 1e). This observation was further supported by scanning electron microscopy data, which confirmed that CD4^+^ EMRA T cells had a significantly higher volume of mitochondria (Figure 1c and d). Together with increased mitochondrial mass we also observed increased levels of the co-transcriptional regulator important for mitochondrial biogenesis; PGC1α, in CD4^+^ EMRA T cells by flow cytometry and qPCR (Figure 1e, extended data figure 1f). Collectively these data positively correlated with the GATA3 expression data and suggested a possible link between GATA3 and mitochondrial biogenesis.

To determine whether the increased mitochondrial mass was of functional importance we used tetramethylrhodamine (TMRE) to measure mitochondrial membrane potential (ΔΨm) in CD4^+^ T cell subsets. Notably, we observed that a greater proportion of CD4^+^ EMRA T cells had hyperpolarised mitochondria compared with the other three subsets (Figure 2a). Hyperpolarised mitochondria have been shown to be a source of reactive oxygen species (ROS) in both macrophages^14^ and CD8^+^ tumour infiltrating lymphocytes^15^. Therefore, we examined intracellular ROS and found that CD4^+^ EMRA T cells also displayed high levels of ROS (Figure 2b). However, although the levels of ROS were significantly higher in the CD4^+^ EMRA T cells compared with all other subsets, the ROS/mitochondria ratio was unchanged (Figure 2b). This led us to postulate that increased mitochondrial mass in CD4^+^ EMRA T cells is a protective mechanism to ensure that CD4^+^ EMRA T cells are able to adequately buffer the excess ROS and prevent further DNA damage.

**Figure 2:**
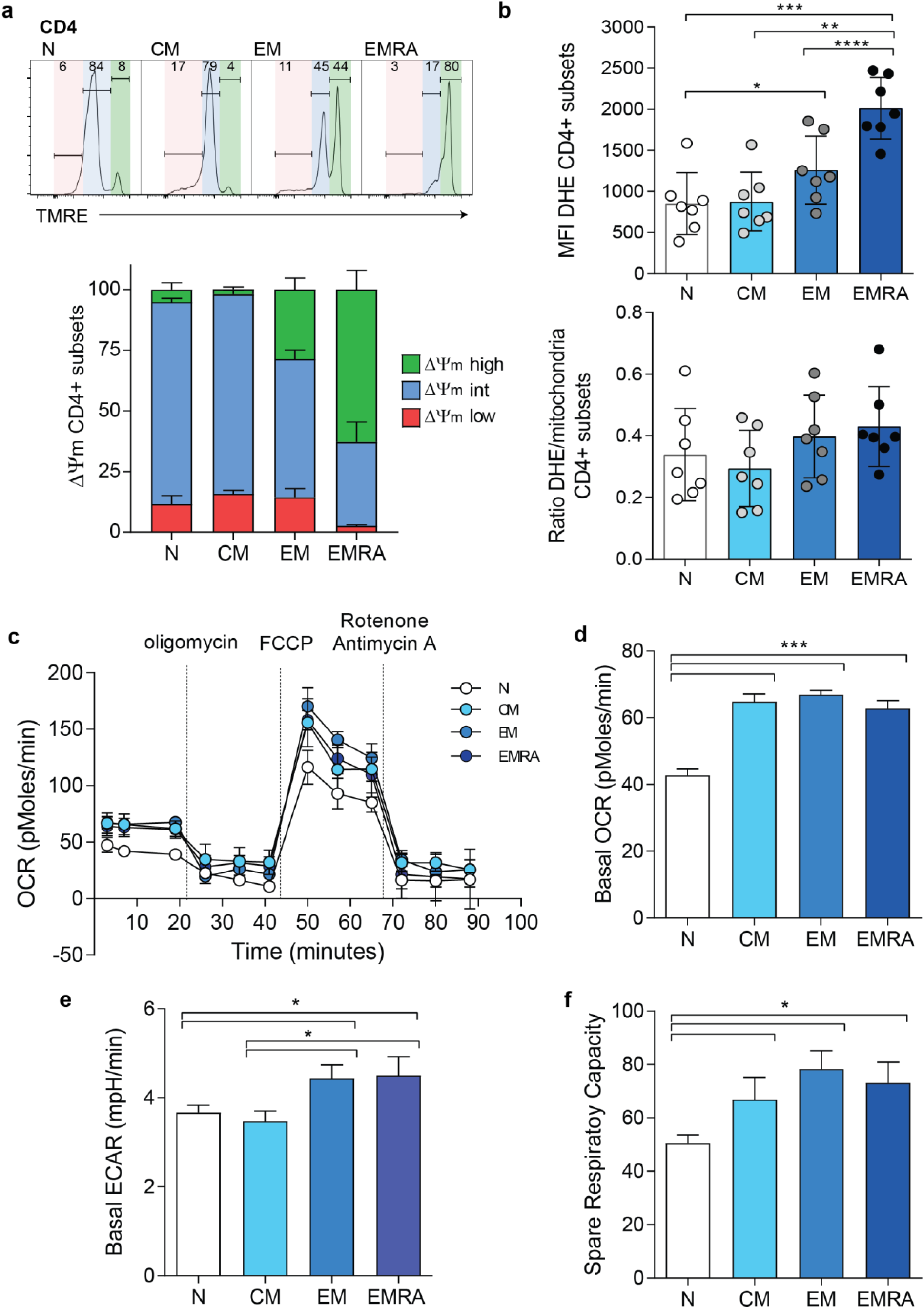
CD4^+^ EMRA T cells have hyperpolarised mitochondria but maintain normal T cell metabolism. (a) Mitochondrial membrane potential (MMP) was determined in CD27/CD45RA defined CD4+ T cell subsets using TMRE (n=10). Low MMP = red, intermediate MMP = blue, high MMP = green. (b) ROS production was obtained using DHE in the CD27/CD45RA defined CD4^+^ T cell subsets (n=7). (c) OCR of the CD4^+^ T cell subsets was measured after 15 minute stimulation with 0.5 μg/ml anti-CD3 and 5 ng/ml IL-2, cells were subjected to a mitochondrial stress test using indicated mitochondrial inhibitors. Data are representative of 4 independent experiments. (d) Basal OCR, (e) Basal ECAR and (f) the SRC of the CD4^+^ T cell subsets after 15 minute stimulation with 0.5 μg/ml anti-CD3 and 5 ng/ml IL-2 (n=4).

We next investigated whether increased mitochondrial mass and hyperpolarised mitochondrial membrane potential in CD4^+^ EMRA T cells led to metabolic reprogramming. To do this we used a mitochondrial stress test to measure the bioenergetics profiles of TCR stimulated CD45RA/CD27 defined CD4^+^ T cell subsets (Figure 2c). The mitochondrial stress test includes the addition of four pharmacological drugs that manipulate oxidative phosphorylation (OXPHOS), which were added in succession to alter the bioenergetics profiles of the mitochondria. Upon stimulation, naïve CD4^+^ T cells had a significantly lower basal oxygen consumption rate (OCR) compared to the memory T cells, however no differences were found amongst the three memory subsets (Figure 2d). The extracellular acidification rate (ECAR), a marker of lactic acid production and glycolysis, was highest in the EM and EMRA CD4^+^ subsets compared to naïve and CM subsets (Figure 2e). Furthermore, the spare respiratory capacity (SRC), a measure of how well mitochondria are potentially able to produce energy under conditions of stress, was also significantly lower in the naïve CD4^+^ T cells but unchanged in the three memory subsets (Figure 2f). CD4^+^ T cells remained metabolically quiescent with no stimulation (Extended data figure 1g). Collectively, these data suggest that despite having hyperpolarised mitochondria CD4^+^ EMRA T cells are able to maintain normal T cell metabolism.

DNA damage can severely affect the function of DNA and plays a major role in age-related diseases and cancer^16^. DNA damage can occur for a number of reasons. For instance, cells which undergo rapid proliferation tend to accumulate DNA damage, as well as many end-stage cells that have undergone replicative senescence in response to telomere dysfunction^17–19^. In support of this we found that DNA damage, determined by an increase in the histone marker for double stranded DNA breaks; ƴH2AX, was significantly higher in the CD4^+^ EMRA T cells when compared with the other three subsets (Figure 3a). To further verify this, activation of the downstream regulator of the DNA damage response; p53 was determined by the phospho-p53/p53 ratio. Indeed, we found that activated p53 increased with differentiation, with the naive CD4^+^ T cells expressing the least and the CD4^+^ EMRA subset expressing the most (Figure 3a). In addition, CD4^+^ EMRA T cells appear to accumulate DNA damage in response to ROS faster than the other memory subsets (Figure 3b).

**Figure 3:**
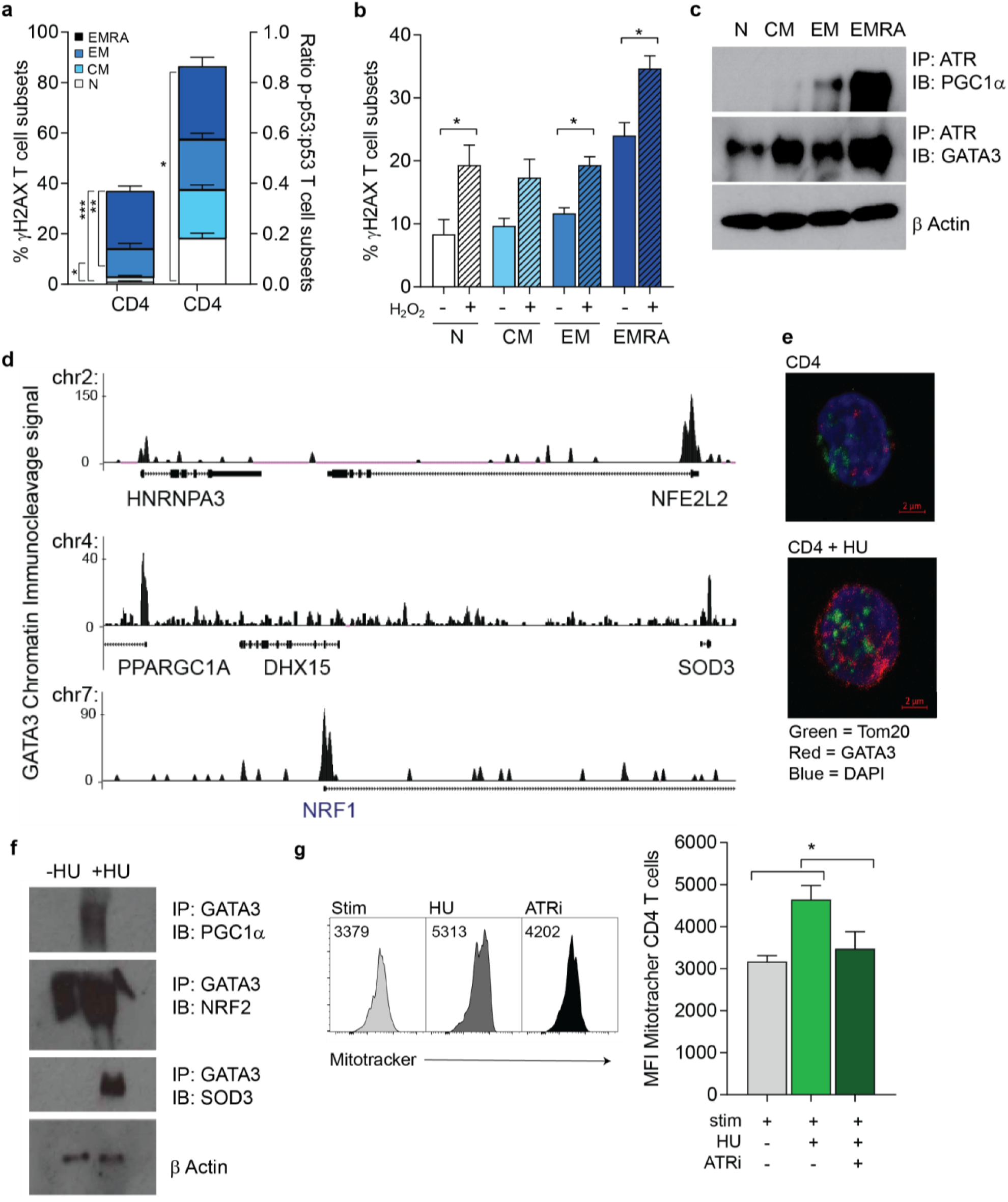
DNA damage recruits a GATA-3-PGC1α complex to induce mitochondria biogenesis in CD4^+^ EMRA T cells. (a) DNA damage in the CD27/CD45RA defined CD4^+^ T cell subsets was determined by assessing the percentage of ƴH2AX positive cells and the p53:p-p53 ratio (n=6). (b) DNA damage in CD4^+^ T cell subsets following the induction of oxidative stress using H_2_O_2_ (n=4). (c) Western blot showing Chk1 immunoprecipitation in the CD27/CD45RA defined CD4^+^ T cell subsets quantifying the presence of GATA3 and PGC1α bound to the Chk1 antibody (n=3). (d) Chromatin immunocleavage showing GATA3 binding to the promoter regions of NRF1, NRF2, SOD3, and PGC1α in naïve CD4^+^ T cells. (e) Tom20 and GATA3 staining in CD4^+^ T cells following treatment with hydroxyurea (n=3). (f) Western blot showing GATA3 immunoprecipitation in CD4^+^ T cells following hydroxyurea treatment quantifying the presence of PGC1α, NRF2 and SOD3 (n=3). (g) Representative flow cytometry histograms and graph of mitochondrial mass measured using mitotracker green following treatment of CD4^+^ T cells with and without hydroxyurea and AZD6738 (n=4).

A possible connection between GATA3 and mitochondria has been previously reported. A yeast two-hybrid screen showed that GATA3 is able to interact with the DDR proteins; ataxia telangiectasia mutated (ATM) and ataxia telangiectasia and Rad3-related (ATR) as well as PGC1α^20^. Immunoprecipitation (IP) assays showed that GATA3 and PGC1α were bound to the ATR in CD4^+^ T cells (Figure 3c). Furthermore, CD4^+^ EMRA T cells had the highest levels of GATA3 and PGC1α bound to the ATR in all three repeats (Extended data figure 2a). We next sort to identify other mitochondrial regulators that may also be upregulated by GATA3 using chromatin immunocleavage.

These data revealed that in resting naïve T cells, GATA3 may be recruited directly or indirectly to the promoter regions of a number of genes including NRF1, NRF2, SOD3, as well as PGC1α (Figure 3d). To mimic DNA damage, we incubated CD4^+^ T cells with hydroxyurea; a DNA replication inhibitor known to induce DNA double strand breaks primarily via the ATR-Chk1 pathway^21,22^. Overnight incubation with hydroxyurea resulted in a significant increase in Tom20, a major outer mitochondrial membrane receptor and therefore mitochondrial marker, and GATA3 using confocal analysis. The increased expression of GATA3 was located in both the cytoplasm and nucleus (Figure 3e, Extended data figure 2b). In addition, flow cytometry revealed a significant increase in phospho-p53, intracellular PGC1α and an increased mitochondrial content, determined by elevated Mitotracker green staining (Extended data figure 2c-e). Furthermore, IP assays showed that treatment with hydroxyurea increased the amount of PGC1α, NRF2 and SOD3 bound to GATA3 (Figure 3f, Extended data figure 2f). Interestingly we found no interaction between NRF1 and GATA3, suggesting that GATA3 may regulate NRF1 at the level of transcription rather than via protein-protein interactions (data not shown).

In order to determine the requirement for ATR activation in regulating mitochondrial biogenesis we used the ATR inhibitor AZD6738 (Ceralasertib). We found that inhibition of the ATR following hydroxyurea treatment prevented the increase in mitochondrial biogenesis, bringing mitochondrial mass down to the level observed with stimulation alone (Figure 3g). We then sought to determine the confirmation of the complex through siRNA knockdown of GATA3 followed by pull down of PGC1α. We found ATR present in both the control and GATA3 siRNA lanes (data not shown) suggesting that while ATR activity is required for complex formation its assembly may not be sequential.

To verify that GATA3 can directly regulate mitochondrial biogenesis and fitness we used small interfering RNA (siRNA) to reduce GATA3 expression in purified human CD4^+^ T cells following activation (Extended data figure 3a-c). Transfection with GATA3 siRNA led to a significant decrease in both the levels of PGC1α and Mitotracker Green staining (Figure 4a, b; Extended data figure 3d), signifying a marked reduction in mitochondrial mass. Confocal microscopy examination of Tom20 (Figure 4c) showed a significant reduction in both mitochondrial mass (Extended data figure 3e) and volume (Extended data figure 3f) in Jurkat cells following transfection with GATA3 siRNA. TMRE staining revealed a dramatic shift from hyperpolarised to hypopolarised mitochondrial membrane potential in the GATA3 siRNA transfected CD4^+^ T cells but not the scrambled siRNA transfected CD4^+^ T cells (Figure 4d). Moreover, transfection with GATA3 siRNA dramatically inhibited the ability of CD4^+^ T cells to perform OXPHOS when compared with CD4^+^ T cells transfected with negative control scrambled siRNA (Figure 4e). Basal OCR and SRC were significantly decreased in the GATA3 siRNA transfected CD4^+^ T cells (Figure 4f). CD8^+^ T cells transfected with GATA3 siRNA showed no difference in their bioenergetic profiles when compared with negative control scrambled siRNA (Extended data figure 3g). Furthermore, we failed to observe a switch to glycolysis with GATA3 knockdown, as ECAR did not significantly increase with the addition of oligomycin (Figure 4g). Additionally, chromatin immunocleavage demonstrated that GATA3 also bound to the promoter regions of hexokinase 1 and 2, together with SLC2A1 suggesting that GATA3 knockdown could interfere with glycolytic metabolism (Extended data figure 4a). A finding that is supported by data showing that GATA3 controls glycolysis through the negative regulation of PPARγ^23^.

**Figure 4:**
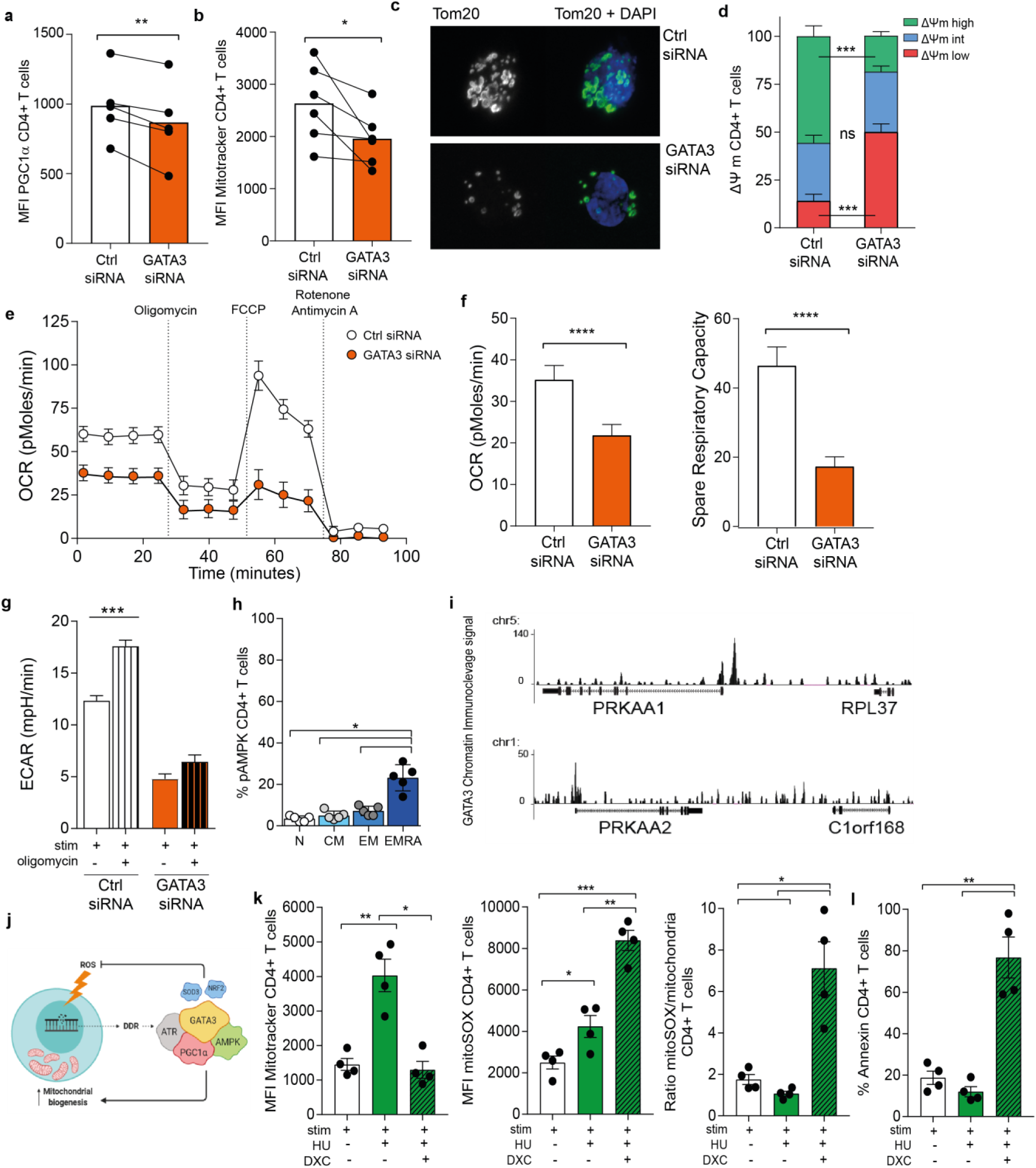
GATA3 and AMPK regulate CD4^+^ T cell metabolism and mitochondrial biogenesis. Graphs showing (a) PGC1α and (b) Mitotracker Green staining in whole CD4^+^ T cells transfected with either control or GATA3 siRNA (n=5 and 6 respectively). (c) Tom20 confocal staining in Jurkat T cells transfected with either control or GATA3 siRNA (n=3). (d) MMP was determined in whole CD4^+^ T cells transfected with either control or GATA3 siRNA using TMRE (*n* = 6). Low MMP = red, intermediate MMP = blue, high MMP = green. (e) OCR of whole CD4+ T cells transfected with either control or GATA3 siRNA was measured after 15 minute stimulation with 0.5 μg/ml anti-CD3 and 5 ng/ml IL-2, cells were subjected to a mitochondrial stress test using indicated mitochondrial inhibitors. Data are representative of 3 independent experiments. (f) Basal OCR and the SRC, together with ECAR following treatment with oligomycin (g) of the CD4^+^ siRNA transfected cells stimulated as described above (n=3). (h) ex-vivo staining of pAMPK (Thr172) in CD27/CD45RA defined CD4+ T cell subsets (n=5). (i) Chromatin immunocleavage showing GATA3 binding to the promoter regions of PRKAA1, PRKAA2 in naïve CD4^+^ T cells. (j) Potential mechanism of GATA3 and AMPK leading to mitochondrial biogenesis during DNA damage. (k) Quantification of mitochondrial mass measured using mitotracker green, and assessment of ROS production using mitoSOX. Together with apoptosis measured by annexin V staining (l) following treatment of CD4^+^ T cells with and without hydroxyurea and doxycycline (n=4).

We then sought to determine how GATA3 regulates the activity of PGC1α. We found that AMPK was highly expressed in the CD4^+^ EMRA subset ex-vivo (Figure 4h, Extended data figure 4b) and expression remained high following stimulation (Extended data figure 4c, d), which correlated to the level of γH2AX observed in CD4^+^ EMRA T cells (Figure 3a). Indeed it has been reported that the ATM plays a role in mitochondrial biogenesis through AMPK activation in response to DNA damage^24^. Furthermore, chromatin immunocleavage revealed that GATA3 can be recruited to the promoter regions of both AMPK1 and 2 (Figure 4i). While c-myc has been shown to play a role in B cell mitochondrial biogenesis^25^, we find no evidence for its involvement in this model. Despite chromatin immuocleavage showing GATA3’s recruitment to the promoter region of c-myc (Extended data figure 4e) we do not see any change in c-myc expression post-transcriptionally following knockdown of GATA3 (Extended data Figure 4f). Mitochondrial biogenesis mediated via c-myc is thought to be regulated by NRF1 gene targets^26^, as we failed to find evidence of NRF1 involvement we believe that NRF1-c-myc do not control mitochondrial biogenesis through DNA damage. Instead, we postulate a feed-forward loop involving GATA3 and AMPK, whereby AMPK phosphorylates PGC1α (Figure 4j).

Finally, we sought to investigate the functional consequence of preventing increased mitochondrial mass in response to DNA damage. We prevented mitochondrial biogenesis using doxycycline, an antibiotic that inhibits mitochondrial protein translation^27^. When doxycycline was given following hydroxyurea treatment, we observed a reduction in mitochondrial content as well as a dramatic increase in ROS production (Figure 4k; extended data figure 4g). Moreover, the decline in mitochondrial content led to a rise in apoptosis, evident by the sharp increase in annexin expression (Figure 4l; extended data figure 4h). Taken together, these data provide evidence that GATA3 modulates mitochondrial biogenesis, cell metabolism, as well as antioxidant response regulating proteins via a nucleus to mitochondria signalling axis to maintain the viability of these cells during DNA damage.

Mechanistically, GATA3 interacts with the DDR protein ATR, activating the PGC1a signalling pathway to promote mitochondrial biogenesis in CD4^+^ T cells, and interacts with antioxidant proteins NRF2 and SOD3, which are important for protecting cells against oxidative damage^28,29^. Importantly, as DDR mechanisms are widely conserved mechanisms, it is likely for this mechanism to not be restricted to the lymphocyte lineage alone. GATA3 is expressed in various cell types outside the haematopoietic system and has been implicated in the tumorigenesis of many cancers, such as luminal breast cancer^30^, neuroblastoma^31^ and endometrial carcinomas^32^.

GATA3-positive breast cancers have been shown to be highly differentiated, with resulting tumours two-fold larger in size than control tumours. In contrast, GATA3-negative breast cancers form poorly differentiated tumours and are highly metastatic^33^. However, despite being associated with low metastatic potential and therefore a more favourable prognosis, GATA3-positive tumours are also associated with resistance to chemotherapy^34^. In light of the present study, we hypothesize that the accumulation of DNA damage generated during chemotherapy recruits GATA3, allowing GATA3-positive tumours to maintain mitochondrial fitness and evade chemotherapy induced apoptosis. However, additional examination of GATA3 and its contribution to mitochondrial biogenesis in cancer is needed to confirm this.

More broadly, our results implicate that transcription factors are likely to serve additional functions after their initial roles in differentiation. This is already the case for the Th1 differentiation marker; T-bet, which is also known to be crucial for long-term memory in CD8^+^ T cells^35,36^ and maturation of NK cells^37^. More recently; STAT5, which has traditionally been associated with cytokine signalling and JAK kinases^38^, has now also been identified as a central regulator of naïve CD4^+^ T cell metabolism^39^. Furthermore, although the GATA proteins have restricted expression patterns to some extent their functions can be interchangeable. For example, GATA1, - 2, and-4 can all activate the expression of GATA3 target genes IL-4 and IL-5 and repress IFNƴ in T cells^40^. Interestingly, GATA4 has previously been shown to become activated in the presence of the ATM and ATR, resulting in the induction of senescence and a senescence associated secretory phenotype (SASP) in human fibroblasts^41^. These functional overlaps suggest that other GATA proteins may also be capable of influencing cell metabolism.

In summary, our data identifies a previously unknown role for GATA3 in response to DNA damage, revealing it as a key mediator of mitochondria biogenesis and cell metabolism. Our findings extend the pleotropic nature of GATA3, demonstrating that more focus should be placed on investigating the role of other transcription factors in later stages of cell development to assess their full range of their abilities. Furthermore, the widespread expression of GATA3 in numerous cancers demonstrate the potential for GATA3-targeted manipulation for clinical interventions.

## Experimental procedures

### Blood sample collection and isolation

Ethics for blood collection were approved by the NRES Committee North East (REC reference: 16/NE/0073) and all subjects provided written informed consent. Heparinized peripheral blood samples were taken from healthy volunteers, average age 41 years ± 5. Healthy volunteers were individuals who had not had an infection or immunisation within the last month, no known immunodeficiency, and had not received any other immunosuppressive medications within the last 6 months. Peripheral blood mononuclear cells (PBMCs) were isolated using Ficoll-Paque (Amersham Biosciences).

### Flow cytometric analysis and cell sorting

Flow cytometric analysis was performed on approximately 1 × 10^6^ PBMCs per sample. Cells were incubated in the dark with the Zombie_NIR live/dead stain (Biolegend) and the following antibodies: CD4 PE-CF594 (RPA-T4) from BD Biosciences, CD45RA BV605 (HI100) and CD27 BV421 (O323) from BioLegend, for 15 minutes at room temperature. For intracellular staining, the following antibodies were used: GATA3 AF488 (16E10A23) and γH2AX AF488 (2F3) from BioLegend, PGC1α (3G6), pAMPK (Thr172, 40H9), p53 Alexa Fluor 647 (1C12) and p-p53 Alexa Fluor 647 (Ser15, 16G8) from Cell Signalling Technology, c-myc (9E10) from Santa Cruz Biotechnology and goat anti-rabbit IgG H&L Alexa Fluor 488 from Abcam.

Detection of GATA3, PGC1α, p53, γH2AX and p-p53 were carried out directly ex vivo. The Foxp3 Transcription Factor Staining Buffer Kit (Biolegend) was used for staining GATA3, p53, γH2AX and p-p53. Following surface staining PBMCs were incubated with the FOXP3 Fix/Perm buffer for 20 minutes in the dark at room temperature followed by incubation with the FOXP3 Perm buffer (10X) for 15 minutes in the dark at room temperature. Finally cells were incubated with the FOXP3 Perm buffer (10X) and appropriate amount of the antibody being used for 30 minutes in the dark and room temperature. The FIX & PERM® Cell Permeablilisation Kit (Life Technologies) was used to enable intracellular staining of PGC1α. Immediately after surface staining, PBMCs were fixed with Reagent A for 15 minutes in the dark at room temperature. Next the cells were permeabilised with Reagent B plus 10% goat serum and recommended volume of primary intracellular antibody. Cells were incubated for 20 minutes in the dark at room temperature followed by a 20 minute incubation with the secondary antibody.

All samples were acquired on a LSR Fortessa (BD Biosciences) equipped with four lasers: 488 nm blue laser, 561 nm yellow green laser, 641 red laser and 405 nm violet laser and data was analysed using FlowJo software.

CD4^+^ T cells were purified using anti-CD4+ conjugated microbeads (Miltenyi Biotec) according to the manufacturer’s instructions. Positively selected CD4+ T cells were labelled with CD27 FITC (O323) and CD45RA APC (HI100) (both from BD Biosciences) and sorted using a FACSAria (BD Biosciences). The purity of CD4 T-cell subsets were assessed by flow cytometry.

### Mitochondrial measurements

For mitochondrial mass, PBMCs were incubated with 100 nM MitoTracker Green FM (Invitrogen(tm)) for 30 minutes at 37°C, 5% CO2. ROS was measured using Dihydroethidium (DHE; ThermoFisher), 5µM MitoSOX was incubated with labelled PBMCs for 20 minutes at 37oC, 5% CO2. Mitochondrial membrane potential (MMP) was investigated using Tetramethylrhodamine (TMRE; ThermoFisher Scientific) 2 µM TMRE was incubated with PBMCs for 30 minutes at 37°C, 5% CO2. Following mitochondrial staining, samples were surface stained and then left unfixed and immediately analysed on an LSR Fortessa (BD Biosciences).

To assess mitochondrial morphology by confocal microscopy, Jurkat cells were plated onto microscope slides then fixed and permed using the FIX & PERM® Cell Permeablilisation Kit (Life Technologies) to enable intracellular labelling. Cells were incubated with Reagent A (Fixation Medium) for 15 minutes at room temperature, then Reagent B (Permeablisaton Medium) plus 10% goat serum and Tom20 (F-10, Santa Cruz) for 20 minutes at room temperature followed incubation with the secondary antibody. Finally, cells were incubated with 4′,6-diamidino-2-phenylindole (DAPI) (ThermoFisher Scientific) for 10 minutes at room temperature in the dark, and then fixed in 2% paraformaldehyde. Samples were imaged on a Zeiss LSM 880 confocal microscope with a X 63 oil-immersion objective lens. Excitation was at 488 nm from an argon-ion laser. Fluorescence detection was in the green; 500 – 570 nm and UV; 405 nm channel. Quantification of image data was performed by measuring the intensity of the 500 – 570 nm channel using ZEN-imaging. Imaris Image Analysis software was used to determine mitochondrial volume through surface 3D rendering of Z-stack images.

### Transmission electron microscopy studies

CD27/CD45RA defined CD4^+^ T cell subsets were isolated and fixed in 2% paraformaldehyde, 1.5% glutaraldehyde in 0.1 m phosphate buffer pH 7.3. They were then osmicated in 1% OsO4 in 0.1M phosphate buffer, dehydrated in a graded ethanol-water series, cleared in propylene oxide and infiltrated with Araldite resin. Ultra-thin sections were cut using a diamond knife, collected on 300 mesh grids, and stained with uranyl acetate and lead citrate. The cells were viewed in a Jeol 1010 transmission electron microscope (Jeol) and imaged using a Gatan Orius CCD camera (Gatan). Mitochondrial volume density (percentage of T cell volume occupied by mitochondria) was determined from EM images using a point-counting method using image J.

### Live cell metabolic assays

The Cell Mito Stress Test was performed on a Seahorse XFe96 Analyser (Agilent) to assess mitochondrial function and OXPHOS. Seahorse sensor cartridges were incubated at 37°C in a non-CO2 incubator overnight with Seahorse XF Calibrant pH 7.4 (Agilent) to rehydrate the sensor probes. The assay was performed in RPMI-1640 buffer without phenol red and carbonate buffer (Sigma-Aldrich) containing 25 mM glucose, 2 mM L-glutamine and 1 mM pyruvate with a pH of 7.4. The stress test was performed using 1 µM Oligomycin, 1.5 µM flurocarbonyl cyanide phenylydrazone (FCCP), 100 nM rotenone, and 1 µM antimycin A (Agilent). Cells were stimulated with 1 µg/µL of anti-CD3 and 5 ng/mL IL-2 15 minutes prior to each metabolic test. Metabolic measurements were calculated according to the manufacturer’s guidelines.

### Immunoprecipitation

For immunoprecipitation experiments the Chk1 or GATA3 antibody was incubated with Red Protein G Affinity beads (Sigma-Aldrich) for 1 hour at 4°C on a shaker to allow the antibody to bind to the beads. Cell lysates were collected in RIPA buffer plus inhibitors (protease: complete mini Roche; 11836153001, and phosphatases: cocktail set II Calbiochem; 524625) and then incubated with the antibody-bead mix overnight at 4°C on a shaker. The following morning, samples were centrifuged, and the supernatant was discarded leaving only the proteins able to bind to Chk1. SDS-polyacrylamide electrophoresis and western blotting of GATA3 and PGC1α was then carried performed as described above. The protein was detected via chemiluminescence, by means of the ECL system (GE Healthcare).

### Hydrogen peroxide experiments

Oxidative stress was induced by the addition of 200 µM H_2_O_2_ for 1 hour. After this time the expression of γH2AX in CD27/CD45RA defined CD4^+^ T cell subsets was assessed by flow cytometry.

### Hydroxyurea experiments

To induce DNA damage, PBMCs were incubated with 400 µM hydroxyurea for 48 hours. Following DNA damage, an immunoprecipitation using the Chk1 antibody was carried out to assess GATA3 and PGC1α binding, apoptosis and mitochondrial properties were examined via flow cytometry. Where indicated 1µM AZD6738 or 50 µM doxycycline was also added to the cell cultures, DMSO was added to control samples.

### Apoptosis experiments

PBMCs were incubated with the desired inhibitors described above, after which time the cells were stained using the Annexin V apoptosis kit (Biolegend) according to the manufacturer’s instructions.

### qPCR of GATA3 and PGC1α

RNA from CD27/CD45RA defined CD4^+^ T cell subsets was isolated using the RNeasy kit (Qiagen) according the manufactures instructions. Transcripts were quantified using the AccuScript high fidelity RT-PCR system (Agilent) according the manufactures instructions. Primer sequences were as follows: GATA3F: GCC CCT CAT TAA GCC CAA G; GATA3R: TTG TGG TCT GAC AGT TCG; PGC1α F: TCT GAG TCT GTA TGG AGT GAC AT; PGC1a R: CCA AGT CGT TCA CAT CTA GTT CA.

### GATA3 siRNA knockdown experiments

GATA-3 and scrambled siRNA (GATA3; 5’-/5Alex647N/UAG GCG AAU CAU UUG UUC AAA-3’, Scrambled; 5’ CCU GUU CUU AAA AUA GUA GGC 3’) were dissolved to a final concentration of 10 µM in electroporation buffer, transferred to an electroporation cuvette and electroporated using program T-023 on an Amaxa Nucleofector (Lonza Bioscience). Cell were then left to recover for 24 to 48 hours in fresh RPMI-1640 media. Primary human CD4+ T cells were stimulated with 1 µg/µL of anti-CD3 and 5 ng/mL IL-2 for 24 hours prior to transfection whereas Jurkat cells were left unstimulated.

### Chromatin immunocleavage

Chromatin Immunocleavage was undertaken using proteinA-micrococcal nuclease (pA-MN)^42^ repurposed to Cut and Run^43^. Naïve primary human T cells were isolated as previously described^44^. Following nuclear isolation, regions of chromatin bound by GATA3 were identified using a mouse anti-GATA3 antibody (HG3-31) Santa Cruz Biotechnology followed by pA-MN. DNA was isolated from eluted chromatin, libraries were prepared using NEBNext® Ultra II DNA Library Prep Master Mix Set and Multiplex Oligos for Illumina® (New England Biolabs). Library quality was assessed using Bioanalyzer 2100 High Sensitivity DNA Gels (Agilent). Libraries were subjected to 50bp single end read sequencing on an HighSeq 2500 (Illumina) in rapid run format and reads were aligned to Human genome hg19 using Bowtie2 (Galaxy v2.2.6)^45^ and visualized within the UCSC genome browser.

### Statistical analysis

GraphPad Prism was used to perform statistical analysis. Statistical significance was evaluated using the paired Student t-test or a repeated-measures ANOVA with the Tukey correction used for post-hoc testing. Differences were considered significant when P was <0.05.

## Acknowledgements

This work was supported by the British Heart Foundation (LAC), a Springboard award from the Academy of Medical Science and the Wellcome trust (ECC, SMH), the Rosetrees Trust (SMH). ANA was funded by the Medical Research Council (MR/P00184X/1). The LSM880 confocal used in these studies was purchased through a Barts and the London Charity grant MGU0293. We would also like to thank the CRUK Flow Cytometry Core Service at Barts Cancer Institute (Core Award<u>C16420</u>/<u>A18066</u>) and the BRC Genomics Platform at National Institute of Health Research (NIHR) Biomedical Research Centre at Guy’s and St. Thomas’ Hospitals, London.

## Data availability

The data that support the findings of this study are available form the corresponding author upon reasonable request.

## Author contributions

LAC, JS and SMH wrote the manuscript, performed the experiments and analysed the data; ECC, EH, AK and PL performed experiments; LELR and JPC assisted with the Tom20 experiments and reviewed the manuscript and ANA reviewed the manuscript.

## Extended data figure legends

**Extended data figure 1:**
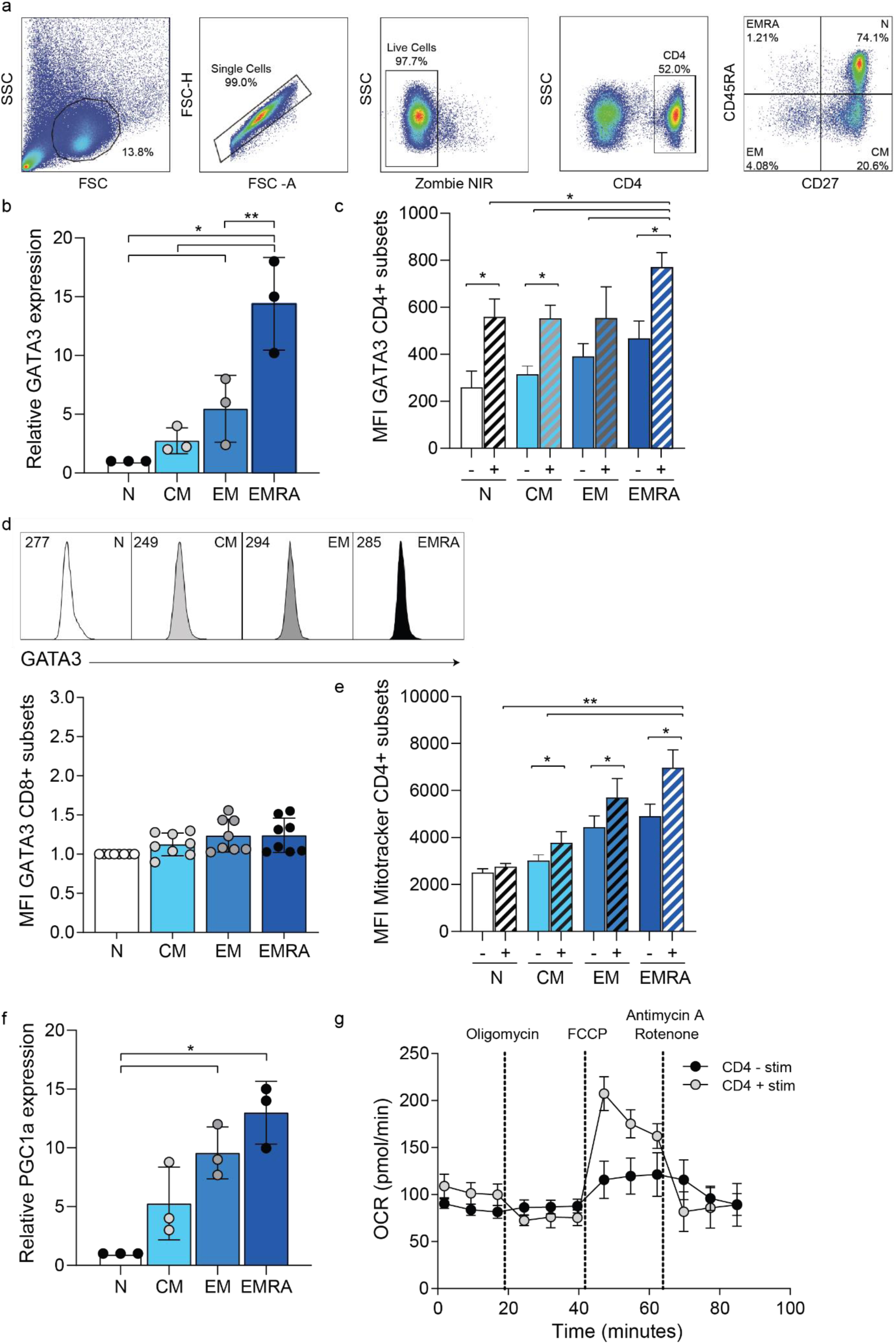
Expression of GATA3 in CD4^+^ T cell subsets. (a) Gating strategy for CD27/CD45RA defined CD4^+^ T cell subsets. From left to right, lymphocytes, single cells, live cells, CD4^+^ T cells and CD27/CD45RA subsets. (b) qPCR data showing the ΔCT relative expression of GATA3 in CD4^+^ T cell subsets (n=3). (c) GATA3 staining in CD27/CD45RA defined CD4^+^ T cell subsets with and without overnight stimulation with 0.5 μg/ml anti-CD3 (n=7) (d) GATA3 expression in unstimulated CD27/CD45RA defined CD8^+^ T cell subsets (n=8). (e) Mitochondrial mass measured by mitotracker green staining in CD27/CD45RA defined CD4^+^ T cell subsets with and without overnight stimulation as described above (n=8). (f) qPCR data showing the ΔCT relative expression of PGC1α in CD4^+^ T cell subsets (n=3). (g) OCR of the CD4^+^ T cells with and without 15 minute stimulation with 0.5 μg/ml anti-CD3 and 5 ng/ml IL-2, cells were subjected to a mitochondrial stress test using indicated mitochondrial inhibitors (n=3).

**Extended data figure 2:**
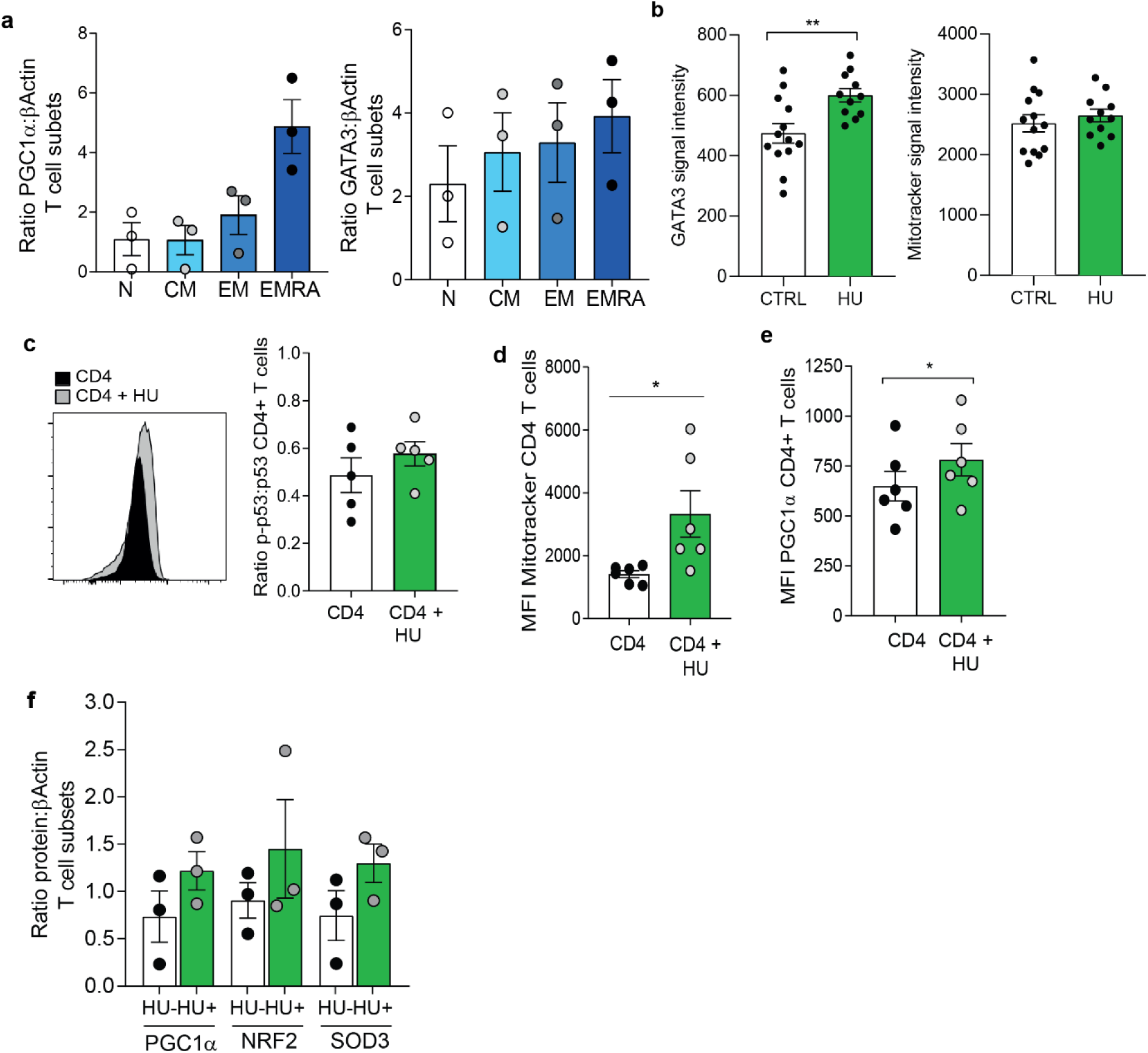
Expression of GATA3 following hydroxyurea treatment. (a) Quantification of PGC1α and GATA3 following immunoprecipitation, shown as the ratio of PGC1α:βActin and GATA3:βActin (n=3). (b) Stain of p-p53 and the p53:p-p53 ratio following the induction of DNA damage by hydroxyurea in whole CD4^+^ T cells. Mitotracker Green staining (c) and PGC1α (d) in CD4^+^ T cells treated with and without hydroxyurea (n=5). (e) Quantification of GATA3 and mitochondrial mass by signal intensity from confocal data (n=3). (f) Quantification of PGC1α NRF1 and SOD3 following immunoprecipitation, shown as the ratio:βActin (n=3).

**Extended data figure 3:**
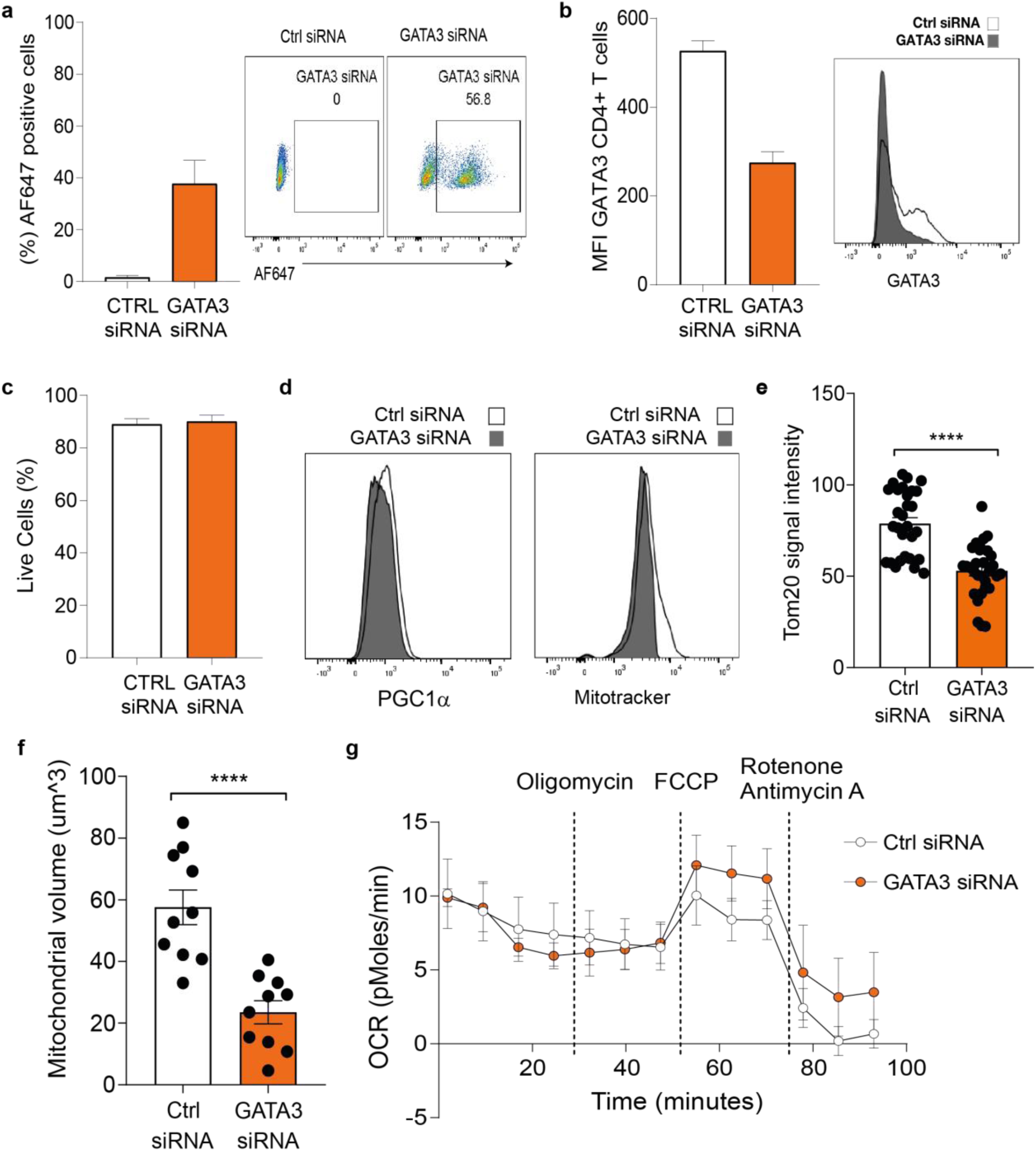
siRNA knockdown of GATA3. (a) GATA3 siRNA transfection efficiency (n=6). (b) GATA3 staining following GATA3 siRNA transfection (n=3). (c) Percentage of live cells determined by Zombie NIR staining following GATA3 siRNA transfection (n=6). (d) Examples of PGC1α expression and mitochondrial mass following siRNA transfection. (e) Mitochondrial mass and (f) mitochondrial volume analysis from confocal microscopy data (n=3). (g) OCR of whole CD8^+^ T cells transfected with either control or GATA3 siRNA was measured after 15 minute stimulation with 0.5 μg/ml anti-CD3 and 5 ng/ml IL-2, cells were subjected to a mitochondrial stress test using indicated mitochondrial inhibitors (n=3).

**Extended data figure 4.**
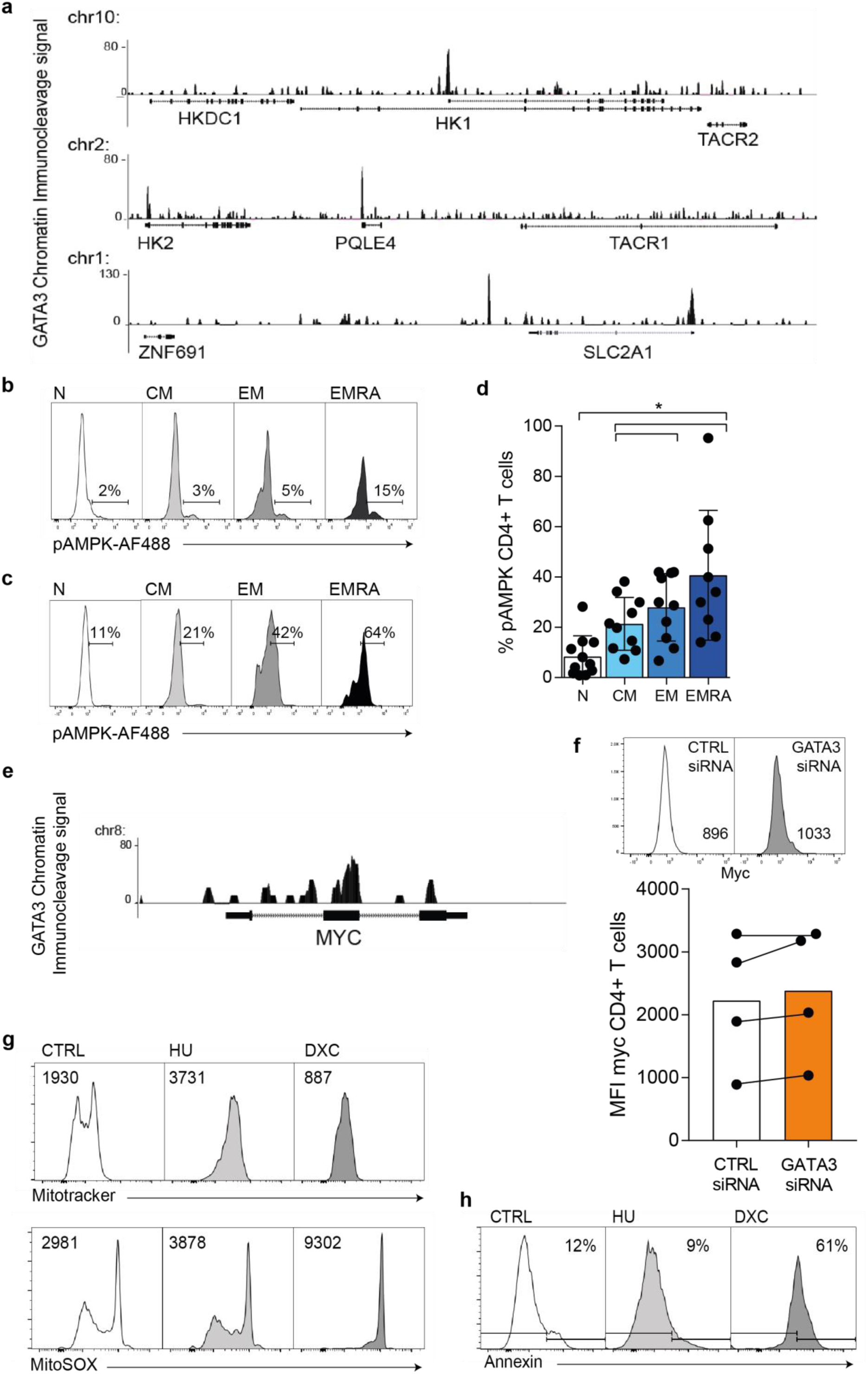
Chromatin immunocleavage and doxycycline treatment. (a) Chromatin immunocleavage showing GATA3 binding to the promoter regions of hexokinase 1/2 and SLC2A1 in naïve CD4^+^ T cells. Representative examples of pAMPK staining ex-vivo (b) and after overnight stimulation (c) with 0.5 ug/ml aCD3. (d) Graph showing pAMPK expression in CD27/CD45RA defined stimulated CD4^+^ T cell subsets (n=9). (e) Chromatin immunocleavage showing GATA3 binding to the promoter regions of c-myc. (f) Data showing c-myc expression in CD4^+^ T cells following transfecting with either a control or GATA3 siRNA (n=4). (g) Representative examples of mitotracker and mitosox staining, together with annexin V staining (h) in CD4^+^ T cells following incubation with hydroxyurea and doxycycline (n=4).

## References

1. Ho, I. C. et al. Human GATA-3: a lineage-restricted transcription factor that regulates the expression of the T cell receptor alpha gene. EMBO J. 10, 1187–92 (2018).

2. Hernández-Hoyos, G., Anderson, M. K., Wang, C., Rothenberg, E. V. & Alberola-Ila, J. GATA-3 expression is controlled by TCR signals and regulates CD4/CD8 differentiation. Immunity 19, 83–94 (2003).

3. Zheng, W. P. & Flavell, R. A. The transcription factor GATA-3 is necessary and sufficient for Th2 cytokine gene expression in CD4 T cells. Cell 89, 587–96 (1997).

4. Pai, S.-Y., Truitt, M. L. & Ho, I.-C. GATA-3 deficiency abrogates the development and maintenance of T helper type 2 cells. Proc. Natl. Acad. Sci. 101, 1993–8 (2004).

5. Kim, P. J. et al. GATA-3 regulates the development and function of invariant NKT cells. J. Immunol. (Baltimore, Md 1950) 177, 6650–9 (2006).

6. Samson, S. I. et al. GATA-3 promotes maturation, IFN-γ production, and liver-specific homing of NK cells. Immunity 19, 701–11 (2003).

7. Tai, T.-S., Pai, S.-Y. & Ho, I.-C. GATA-3 Regulates the Homeostasis and Activation of CD8+ T Cells. J. Immunol. 190, 428–37 (2012).

8. Das, A. et al. Effector/memory CD4 T cells making either Th1 or Th2 cytokines commonly co-express T-bet and GATA-3. PLoS One 12, e0186932 (2017).

9. Sakata-Kaneko, S., Wakatsuki, Y., Matsunaga, Y., Usui, T. & Kita, T. Altered Th1/Th2 commitment in human CD4+ T cells with ageing. Clin. Exp. Immunol. 120, 267–73 (2000).

10. Koch, S. et al. Multiparameter flow cytometric analysis of CD4 and CD8 T cell subsets in young and old people. Immun. Ageing 5, 6 (2008).

11. Henson, S. M., Riddell, N. E. & Akbar, A. N. Properties of end-stage human T cells defined by CD45RA re-expression. Curr. Opin. Immunol. 24, 476–81 (2012).

12. Di Mitri, D. et al. Reversible Senescence in Human CD4+CD45RA+CD27-Memory T Cells. J. Immunol. 187, 2093–100 (2011).

13. Henson, S. M. et al. P38 signaling inhibits mTORC1-independent autophagy in senescent human CD8+ T cells. J. Clin. Invest. 124, 4004–16 (2014).

14. Mills, E. L. et al. Repurposing mitochondria from ATP production to ROS generation drives a pro-inflammatory phenotype in macrophages that depends on succinate oxidation by complex II. Cell 167, 457–470 (2016).

15. Siska, P. J. et al. Mitochondrial dysregulation and glycolytic insufficiency functionally impair CD8 T cells infiltrating human renal cell carcinoma. JCI Insight 2, 93411 (2017).

16. Hoeijmakers, J. H. J. DNA Damage, Aging, and Cancer. N. Engl. J. Med. 361, 1475–1485 (2009).

17. Hayflick, L. & Moorhead, P. S. The serial cultivation of human diploid cell strains. Exp. Cell Res. 25, 585–621 (1961).

18. Harley, C. B., Futcher, A. B. & Greider, C. W. Telomeres shorten during ageing of human fibroblasts. Nature 345, 458–60 (1990).

19. Richter, T. & Zglinicki, T. von. A continuous correlation between oxidative stress and telomere shortening in fibroblasts. Exp. Gerontol. 42, 1039–42 (2007).

20. Kim, M. A. et al. Identification of novel substrates for human checkpoint kinase Chk1 and Chk2 through genome-wide screening using a consensus Chk phosphorylation motif. Exp. Mol. Med. 39, 205–12 (2007).

21. Bianchi, V., Pontis, E. & Reichard, P. Changes of deoxyribonucleoside triphosphate pools induced by hydroxyurea and their relation to DNA synthesis. J. Biol. Chem. 261, 16037–16042 (1986).

22. Timson, J. Hydroxyurea. Mutat. Res. 32, 115–132 (1975).

23. Lee, J. E. & Ge, K. Transcriptional and epigenetic regulation of PPARγ expression during adipogenesis. Cell Biosci 4, 29 (2014).

24. Fu, X., Wan, S., Lyu, Y. L., Liu, L. F. & Qi, H. Etoposide induces ATM-dependent mitochondrial biogenesis through AMPK activation. PLoS One 3, e2009 (2008).

25. Li, F. et al. Myc Stimulates Nuclearly Encoded Mitochondrial Genes and Mitochondrial Biogenesis Graduate Programs in Human Genetics and Molecular Biology. Mol. Cell. Biol. 25, 6225–6234 (2005).

26. Kim, J., Lee, J. H. & Iyer, V. R. Global identification of Myc target genes reveals its direct role in mitochondrial biogenesis and its E-box usage in vivo. PLoS One 3, e1798 (2008).

27. Lamb, R. et al. Antibiotics that target mitochondria effectively eradicate cancer stem cells, across multiple tumor types: Treating cancer like an infectious disease. Oncotarget 6, 4569–84 (2015).

28. Nguyen, T., Nioi, P. & Pickett, C. B. The Nrf2-antioxidant response element signaling pathway and its activation by oxidative stress. J. Biol. Chem 284, 13291–5 (2009).

29. Singh, B. & Bhat, H. K. Superoxide dismutase 3 is induced by antioxidants, inhibits oxidative DNA damage and is associated with inhibition of estrogen-induced breast cancer. Carcinogenesis 33, 2601–10 (2012).

30. Mehra, R. et al. Identification of GATA3 as a breast cancer prognostic marker by global gene expression meta-analysis. Cancer Res. 65, 11259–11264 (2005).

31. Peng, H. et al. Essential role of GATA3 in regulation of differentiation and cell proliferation in SK-N-SH neuroblastoma cells. Mol. Med. Rep. 11, 881–886 (2015).

32. Engelsen, I. B., Stefansson, I. M., Akslen, L. A. & Salvesen, H. B. GATA3 expression in estrogen receptor α-negative endometrial carcinomas identifies aggressive tumors with high proliferation and poor patient survival. Am. J. Obstet. Gynecol. 199, 543.e1– 7 (2008).

33. Kouros-Mehr, H. et al. GATA-3 Links Tumor Differentiation and Dissemination in a Luminal Breast Cancer Model. Cancer Cell 13, 141–52 (2008).

34. Tominaga, N. et al. Clinicopathological analysis of GATA3-positive breast cancers with special reference to response to neoadjuvant chemotherapy. Ann. Oncol. 23, 3051–7 (2012).

35. Juedes, A., Glimcher, L. H., Szabo, S. J., von Herrath, M. & Sullivan, B. M. Antigen-driven effector CD8 T cell function regulated by T-bet. Proc. Natl. Acad. Sci. 100, 15818–23 (2003).

36. Intlekofer, A. M. et al. Effector and memory CD8+ T cell fate coupled by T-bet and eomesodermin. Nat. Immunol. 6, 1236–44 (2005).

37. Townsend, M. J. et al. T-bet regulates the terminal maturation and homeostasis of NK and Vα14i NKT cells. Immunity 20, 477–94 (2004).

38. Beyer, T. et al. Integrating signals from the T-cell receptor and the interleukin-2 receptor. PLoS Comput. Biol. 7, e1002121 (2011).

39. Jones, N. et al. Akt and STAT5 mediate naïve human CD4+ T-cell early metabolic response to TCR stimulation. Nat. Commun. 10, 2042 (2019).

40. Ranganath, S. & Murphy, K. M. Structure and Specificity of GATA Proteins in Th2 Development. Mol. Cell. Biol. 21, 2716–25 (2002).

41. Kang, C. et al. The DNA damage response induces inflammation and senescence by inhibiting autophagy of GATA4. Science. 349, aaa5612 (2015).

42. Schmid, M., Durussel, T. & Laemmli, U. K. ChIC and ChEC: Genomic mapping of chromatin proteins. Mol. Cell 16, 147–57 (2004).

43. Skene, P. J. & Henikoff, S. A simple method for generating highresolution maps of genome-wide protein binding. Elife 4, e09225 (2015).

44. Cousins, D. J., Lee, T. H. & Staynov, D. Z. Cytokine Coexpression During Human Th1/Th2 Cell Differentiation: Direct Evidence for Coordinated Expression of Th2 Cytokines. J. Immunol. 169, 2498–2506 (2002).

45. Afgan, E. et al. The Galaxy platform for accessible, reproducible and collaborative biomedical analyses: 2016 update. Nucleic Acids Res. 46, W537–W544 (2016).

